# Machine learning–driven decoding of maternal immune signatures in repeated pregnancy loss

**DOI:** 10.1101/2025.08.29.673197

**Authors:** Tae Lyun Ko, Jaesub Park, Dongju Leem, Junho Kim, Jae won Han, Jin Sol Park, Sung Ki Lee, Hyojung Paik

**Affiliations:** Center for Biomedical Computing, Korea Institute of Science and Technology Information (KISTI), and Department of Data and HPC Science, Korea National University of Science and Technology (UST), Daejeon 34141, Republic of Korea; Department of Bio and Brain Engineering, Korea Advanced Institute of Science and Technology, Daejeon 34141, Republic of Korea, and Cambridge Stem Cell Institute, University of Cambridge, Cambridge, UK; Department of Obstetrics and Gynecology, College of Medicine, Myunggok Medical Research Institute, Konyang University, Daejeon 158, Republic of Korea; Department of Biological Sciences, Sungkyunkwan University, Suwon, 16419, Republic of Korea

**Author notes:** These authors are equally contributed (Tae Lyun Ko, Jaesub Park).

## Abstract

**Background:** Repeated pregnancy loss (RPL) is a multifactorial condition in which the underlying immunological mechanisms remain incompletely understood. Although immune tolerance at the maternal–fetal interface is critical for successful pregnancy, the immune disruptions that contribute to RPL, independent of fetal aneuploidy, remain poorly characterized.

**Methods:** We performed single-cell RNA sequencing of decidual tissues from RPL patients and gestational age–matched controls. We employed single-cell transcriptomics coupled with genotype-based origin analysis to dissect immune dysregulation in RPL. To identify cell type-specific RPL signatures, we mutually applied a supervised machine learning model and foundation model-based analysis, followed by pathway enrichment and network analysis. In addition, we prioritized drug repurposing candidates based on the drug response data, which reversed the RPL-associated transcriptional profiles.

**Results:** In normal pregnancy, fetal immune cells were nearly absent but increased in RPL, whereas fetal trophoblasts were reduced, suggesting hindered placental development despite the absence of chromosomal abnormalities in fetal cells. Interestingly, by showing a strong common rejection module score, RPL immune cells resemble the transcriptional signatures of acute transplant rejection, indicating the contribution of the maternal immune system. Our machine learning and transformer-based models mutually identified T-cell–derived transcriptomic signatures that distinguished RPL immune cells. By examining biological confounding factors, including fetal-origin signatures, we prioritized CXCR4 and JUN in maternal T cells as RPL-associated signatures. For clinical application, we explored reversed transcriptional signatures from drug response data, and three compounds were highlighted as candidates for drug repurposing.

**Conclusions:** Together, our approach identifies maternal immune signatures such as those of CXCR4 and JUN and links them to potential drug repurposing candidates, thereby providing both mechanistic insights and therapeutic opportunities for RPL.

## Background

Repeated pregnancy loss (RPL) is defined as the loss of two or more consecutive pregnancies before 20 weeks of gestation(1) and represents a major area of investigation for understanding the underlying mechanisms during the early stages of fetal development and fertility. Previous studies have suggested that immune tolerance between the mother and fetus is an essential prerequisite for maintaining a successful pregnancy.(2) Disruption of this immunological tolerance has been proposed as an usual suspect of RPL.(3) However, the single-cell-level understanding of the immune coordination between semiallogeneic cells, such as feto-maternal cells, remains poorly understood with respect to their cellular origin. To address this gap, we applied a machine learning-based approach to single-cell transcriptional profiles from RPL samples, aiming to prioritize cell types—distinguished by their origin—that are most strongly associated with RPL.

RPL is a clinically heterogeneous condition with multiple potential causes, including anatomical defects, endocrine disorders, thrombophilia, and immunological dysfunction.(4) Previous attempts have focused on fetal chromosomal abnormalities, particularly aneuploidy, as potential contributors to RPL. For instance, Stephenson et al. conducted cytogenetic analyses of miscarriage tissues from RPL patients and identified chromosomal anomalies in a subset of cases. Although such aneuploidy analyses improve our conceptual understanding of early fetal development, therapeutic application in aneuploid pregnancies remains limited.

Given these limitations, we focused on maternal etiology, which may contribute to pregnancy loss in the absence of identifiable chromosomal errors. A wide range of maternal contributions have been implicated in RPL, including uterine anomalies, hormonal imbalances, thrombophilic conditions, and immune dysregulation.(5) Among these, immune-related mechanisms have drawn increasing amounts of evidence, with studies suggesting that failures in maternal–fetal immune tolerance may impair implantation and pregnancy maintenance. For example, Deshmukh et al. reported that disruption of immune tolerance at the maternal–fetal interface may contribute to RPL and other gestational complications.(6) However, many of these findings are based on peripheral blood profiling or bulk tissue analysis, which may indirectly reflect immune dynamics at the maternal–fetal interface. Consequently, a more refined understanding of tissue-specific, cell-level immune alterations in euploid RPL remains elusive.

Single-cell RNA sequencing (scRNA-seq) has emerged as a powerful tool for resolving immune heterogeneity directly at the maternal–fetal interface. scRNA-seq enables detailed mapping of cellular heterogeneity and immune architecture in the decidua, providing insight into maternal–fetal communication during early pregnancy. For instance, Vento-Tormo et al. reconstructed the cellular and molecular landscape of the maternal–fetal interface by analyzing thousands of decidual cells from first-trimester pregnancies in normals.(7) Similarly, Suryawanshi et al. profiled the immune and nonimmune cell types in both the placenta and decidua, highlighting the dynamic interactions involved in the early gestation of normals.(8) These studies have expanded our understanding of the maternal–fetal interface at single-cell resolution. However, the complexity of such data highlights the need for approaches that can detect subtle, cell-type–specific transcriptional changes essential for understanding RPL.

Given this analytical complexity, single-cell datasets benefit from advanced computational techniques capable of capturing subtle transcriptional variation. Recent advances in machine learning and artificial intelligence (AI) have provided new opportunities for analyzing high-dimensional single-cell datasets. These methods enable the detection of complex, multivariate transcriptomic patterns that may not be evident through conventional differential expression analysis.(9) In particular, machine learning models such as decision trees and neural networks can capture nonlinear relationships and classify cells based on subtle combinations of gene expression features.(10) More recently, transformer-based architectures in AI have been adapted for single-cell data, where attention mechanisms can highlight context-dependent relationships between genes across cells.(11)

To address these issues, we constructed a single-cell atlas of the human decidua by comparing tissues from normal pregnancies and RPL cases. We tested the hypothesis that RPL stems from a failure of maternal immune tolerance by comparing its cellular signatures to those from acute organ transplant rejection. We then employed machine learning and AI to identify key immune populations and characterize transcriptomic differences that would be related to maternal immune tolerance in RPL. This integrative framework enables single-cell–level dissection of immune dysregulation in the decidua and reveals immune-related mechanisms that may contribute to RPL.

## Methods

### Sample collection

Decidual tissue (transformed endometrium during pregnancy) samples were collected from seven pregnant individuals, including three with normal pregnancies and four with RPL. All participants provided written informed consent, and the study was approved by the Institutional Review Board (IRB) of Konyang University Hospital (IRB approval number: KYUH 2022-08-009-005). In both groups, decidual tissue and peripheral blood were collected via dilation and curettage (D&C). In the normal group, samples were obtained during the elective termination of pregnancy, whereas in the RPL group, two resulted from spontaneous miscarriage, and two were due to elective termination. Chromosomal karyotyping revealed a 46, XX karyotype in both miscarriage cases and one elective termination case, whereas the karyotype of the other elective termination case could not be determined. All the samples were collected under sterile conditions and immediately processed for further analysis, including single-cell RNA sequencing (scRNA-seq).

### Analysis of droplet-based single-cell RNA sequencing data

Single-cell suspensions were prepared through standard tissue dissociation and cell processing procedures. Single-cell RNA libraries were constructed using the 10X Genomics Chromium Platform in accordance with the manufacturer’s instructions. Libraries were sequenced using both the Illumina HiSeq-x and NovaSeq 6000 platforms. The raw sequencing reads were aligned to the reference genome (GRCh38) using Cell Ranger (v5.0.0) to generate gene‒cell count matrices. Ambient RNA noise was minimized using SoupX,(12) and potential doublets were removed with Scrublet.(13) Preprocessing and quality control were conducted in Scanpy (v1.10.1), filtering out cells with mitochondrial gene expression exceeding 20%, fewer than 200 detected genes, fewer than 1,000 UMI counts, or more than 7,000 total genes per cell. The merged data across all samples included a total of 62,588 cells for downstream analysis. The merged data underwent normalization, scaling, and the identification of highly variable genes, followed by dimensionality reduction using PCA and cluster visualization with UMAP.(14) Batch effects were corrected using Harmony(15) after the clustering results were compared with those of BBKNN.(16) Cell types were identified using a logistic regression modeling approach implemented in sceleto2 (https://pypi.org/project/sceleto2/), in conjunction with CellTypist,(17) based on reference labels for cells in the human decidua during the first trimester.(7)

### Identification of fetal and maternal cells

Given the nature of the samples, which were obtained from the decidua of pregnant women following pregnancy termination, the tissues contain a mixture of maternal and fetal cells. Therefore, to accurately determine whether the acquired cells originated from the mother or the fetus, we employed two independent analytical methods for cross-validation. This approach enabled us to reliably identify the origin of each cell. We first utilized Souporcell(18) to cluster the cells based on their genotypes, dividing them into two genetically distinct clusters. To verify the origin of each cluster, we subsequently compared the genotype profiles generated by Souporcell with the genetic variant data obtained from maternal peripheral blood mononuclear cells (PBMCs) using the Illumina GSA v3 genotyping array. The single-nucleotide polymorphisms (SNPs) observed in the two clusters were compared with the maternal peripheral blood mononuclear cell (PBMC) genomic data to assess their similarity. The cluster with higher similarity was determined to consist of maternal cells and was defined as the “maternal cell cluster,” whereas the cluster with lower similarity was considered to consist of fetal cells. To achieve this goal, an average of 927.3 ± 435.3 alleles per sample were examined, allowing us to genetically distinguish the two clusters and identify the origin of each cell.

### Identification of single cells with aneuploidy from scRNA-seq data

Chromosomal aneuploidy was investigated in each cell of the scRNA-seq dataset using the method previously described by Starostik et al.(19) Briefly, chromosomal aneuploidy can be inferred by integrating two types of information: chromosome-wide differential expression of dosage-associated genes and allelic imbalance at heterozygous SNP sites. For each single cell, the expression levels of dosage-associated genes were quantified using scploid, which provides a predefined set of dosage-associated genes along with benchmarking data from mosaic aneuploid mouse embryos.(20) Allelic imbalance was estimated per chromosome by measuring allelic read counts at each heterozygous SNP site.

To apply this method to our scRNA-seq dataset, we first generated aligned BAM files for every single cell using a custom python script that split the BAM files by cell barcode. Based on cell cluster information obtained from Souporcell (see above), maternal and fetal pseudobulk BAM files were generated by merging cells using SAMtools (version 1.3.1).(21) Heterozygous SNPs were identified from the pseudobulk using GATK HaplotypeCaller.(22) Allelic read counts were then measured for each heterozygous SNP site using GATK ASEReadCounter for every cell. Owing to the sparsity of 10X Chromium scRNA-seq data, we included only SNP sites with at least three mapped reads per cell. Based on the measured read counts, we computed the allelic ratio for each chromosome–cell pair by dividing the sum of minor allele reads by the total number of mapped reads. Aneuploidy was determined if the scploid-derived p value was ≤ 0.05 and if the allelic ratio was within the top 10% across all chromosome–cell pairs.

### Calculation of the common rejection module score

The common rejection module (CRM) score is calculated using the expression levels of 11 predefined CRM genes: BASP1, CD6, CXCL10, CXCL9, INPP5D, ISG20, LCK, NKG7, PSMB9, RUNX3, and TAP1.(23) These genes were identified through meta-analyses as being consistently overexpressed during acute rejection (AR) across multiple transplanted organs. The score is derived by averaging the expression values of these CRM genes and subtracting the average expression of a dynamically selected set of control genes. This process ensures that the score reflects the relative activity of rejection-associated genes while accounting for background transcriptional noise. The calculations were performed using the Scanpy package.

### Differential gene expression analysis

Differential gene expression analysis was performed using the Wilcoxon rank-sum test as implemented in scanpy.tl.rank_genes_groups, with Benjamini–Hochberg correction applied for multiple testing. All analyses were conducted on log-normalized gene expression values. Genes were considered differentially expressed if the adjusted *p* value was less than 0.05 and if the absolute log fold change exceeded 1. Comparisons were made between biological conditions of interest, including maternal versus fetal origin and RPL versus normal samples, depending on the analysis context. For a analysis focused on the aneuploid cell subset, we additionally employed the DESeq2 package in R(24). This approach was used to compare normal versus RPL conditions exclusively within the fetal aneuploid cell population.

### Machine learning model

We utilized devCellPy, a hierarchical machine learning framework built on XGBoost, to identify key features associated with the Norm and RPL conditions.(25) The classification model included two stages. In the first level, broad cell types were identified, and in the second level, immune cell types were distinguished based on the Norm and RPL conditions. We assessed the contribution of individual genes in the classification model by leveraging SHAP (Shapley Additive Explanations) analysis, which quantified the impact of each feature on the model’s predictions by attributing a specific importance value to each gene.

### Validation of prioritized RPL-associated genes using a transformer model of AI

To validate the SHAP-derived gene signatures for RPL immune signature identification, we employed scGPT, a transformer-based large language model specialized for single-cell RNA-seq data.(26) For each of the three immune cell types (T cells, granulocytes, and NK CD16+ cells), we selected the top *n* genes ranked by SHAP values from both classification levels of devCellPy. Using the pretrained scGPT_human model, we fine-tuned scGPT exclusively on these selected genes. The fine-tuned models were subsequently used to classify cells into six categories: Tcell_Normal, Tcell_RPL, Granulocyte_Normal, Granulocyte_RPL, NK_CD16⁺_Normal, and NK_CD16⁺_RPL. This validation step ensured that the SHAP-ranked genes retained their predictive power across a different model architecture, reinforcing the robustness of the identified gene signatures to detect RPL-associated immune signatures.

### Drug repurposing

Drug repurposing analysis was performed using ASGARD.(27) As input, we used the signature gene list derived from the integration of patient and cell type information from the previous step. Therefore, patient identity and cell type were not separately used in this step. To ensure adequate coverage across drugs, we selected six tissues (lung, prostate, skin, breast, kidney, and large intestine) for which experimental data were available for at least 700 compounds. Drugs were selected if they were predicted to have significant effects (FDR < 0.05) in at least three of these tissues. Unless otherwise specified, all procedures followed the original ASGARD pipeline.

### Protein target and interaction data

Drug–target interactions were extracted from the DrugBank database.(28) Protein–protein interaction (PPI) data were obtained from the STRING database.(29) To ensure high-confidence interactions, we applied a confidence score threshold of 200 based on the STRING-provided scores. The optimal threshold was determined by varying the cutoff in increments of 100 and selecting the highest value that retained interactions involving all signature proteins and drug targets. Network construction and centrality analyses were performed using the NetworkX library (3.4.2) in Python.(30)

## Results

### Patient statistics

First-trimester decidual biopsy specimens were obtained from women undergoing elective termination of uncomplicated pregnancies (Norm; n = 3; mean gestational age = 8.7 ± 1.5 weeks) and from patients with RPL (n = 4; mean gestational age = 10.6 ± 2.5 weeks) (Table S1). Decidual tissue and matched peripheral blood samples were collected from both Norm and RPL participants. Peripheral blood mononuclear cells (PBMCs) were isolated from peripheral blood by density-gradient centrifugation. In the Norm group, samples were obtained after elective termination; in the RPL group, samples were collected after spontaneous miscarriage. Maternal PBMCs from all participants were genotyped using the Illumina GSA v3 array to inform maternal versus fetal cell-origin assignment.

Fetal chromosomal abnormalities can alter maternal immune responses and may confound our analysis,(31) and immunomodulatory therapies also have limited applicability in such cases. To mitigate this confounding, we performed chromosomal karyotyping of decidual tissues prior to single-cell RNA sequencing. One RPL case with chromosome 9 duplication was identified and excluded from single-cell RNA sequencing.

### Single-cell transcriptomic profiling reveals altered maternal–fetal cell composition in the decidua of RPL

To comprehensively characterize the cellular composition and transcriptional landscape of the decidua during early pregnancy, we isolated single cells from decidual samples and performed scRNA-seq (see Methods). After standard preprocessing and quality control, one Norm case was excluded because of low sequencing quality, and a total of 62,588 cells from two Norm and three RPL cases were retained for analysis (see Methods).

To determine the origins of cells as either maternal or fetal cells, we applied a dual-verification approach, as we presented in our previous attempts.(32) In summary, we identified maternal and fetal origins by clustering cells based on allele variations using Souporcell(18) and subsequently validated the classifications by comparison with genotypes derived from maternal PBMCs (see the Methods.) More than 900 alleles (mean of 927.3 ± 435.3 per sample) were examined to support the assignment of cellular origin. After removing a subset of cells that lacked sufficient allelic information for confident classification as either maternal or fetal, we retained 57,976 cells for further analysis. Among these, 35,027 (60.4%) cells were from RPL samples, and 22,949 (39.6%) were from Norm samples; 41,517 (71.6%) were classified as maternal, and 16,459 (28.4%) were classified as fetal.

Next, we performed unsupervised clustering and cell-type annotation to resolve 32 distinct decidual cell clusters across immune and nonimmune compartments and visualized the data with UMAP (Uniform Manifold Approximation and Projection),(14) colored by cell type, clinical condition (Norm vs. RPL), and inferred origin (maternal vs. fetal) (Fig. 1A–C). Nonimmune cells included decidual stromal cells (dS1, dS2, and dS3), syncytiotrophoblasts (SCT), villous cytotrophoblasts (VCT), extravillous trophoblasts (EVT), endothelial cells (Endo (m), Endo (f), and Endo L), and epithelial glandular cells (Epi1 and Epi2). Immune cells included granulocytes, decidual natural killer cells (dNK1, dNK2, and dNK3), CD16-positive and CD16-negative natural killer cells (NK CD16+ and NK CD16-), decidual macrophages (dM1, dM2, and dM3), dendritic cells (DC1 and DC2), Hofbauer cells (HB), and innate lymphoid cells (ILC3). The inferred cell origins were successfully aligned with known biology, lending support to the robustness of our genotype-guided classification. Trophoblast cells— including EVT, SCT, and VCT cells—were overwhelmingly classified as fetal in origin. For instance, among 3,469 EVT cells, only 5 were assigned a maternal origin, and similar fetal dominance was observed in the SCT and VCT clusters. Likewise, Hofbauer (HB) cells, known as fetal macrophages, were predominantly labeled as fetal (Fig. 1E). These observations are consistent with established knowledge regarding the fetal origin of trophoblast and HB cells, reinforcing the validity of our classification approach.

**Figure 1.**
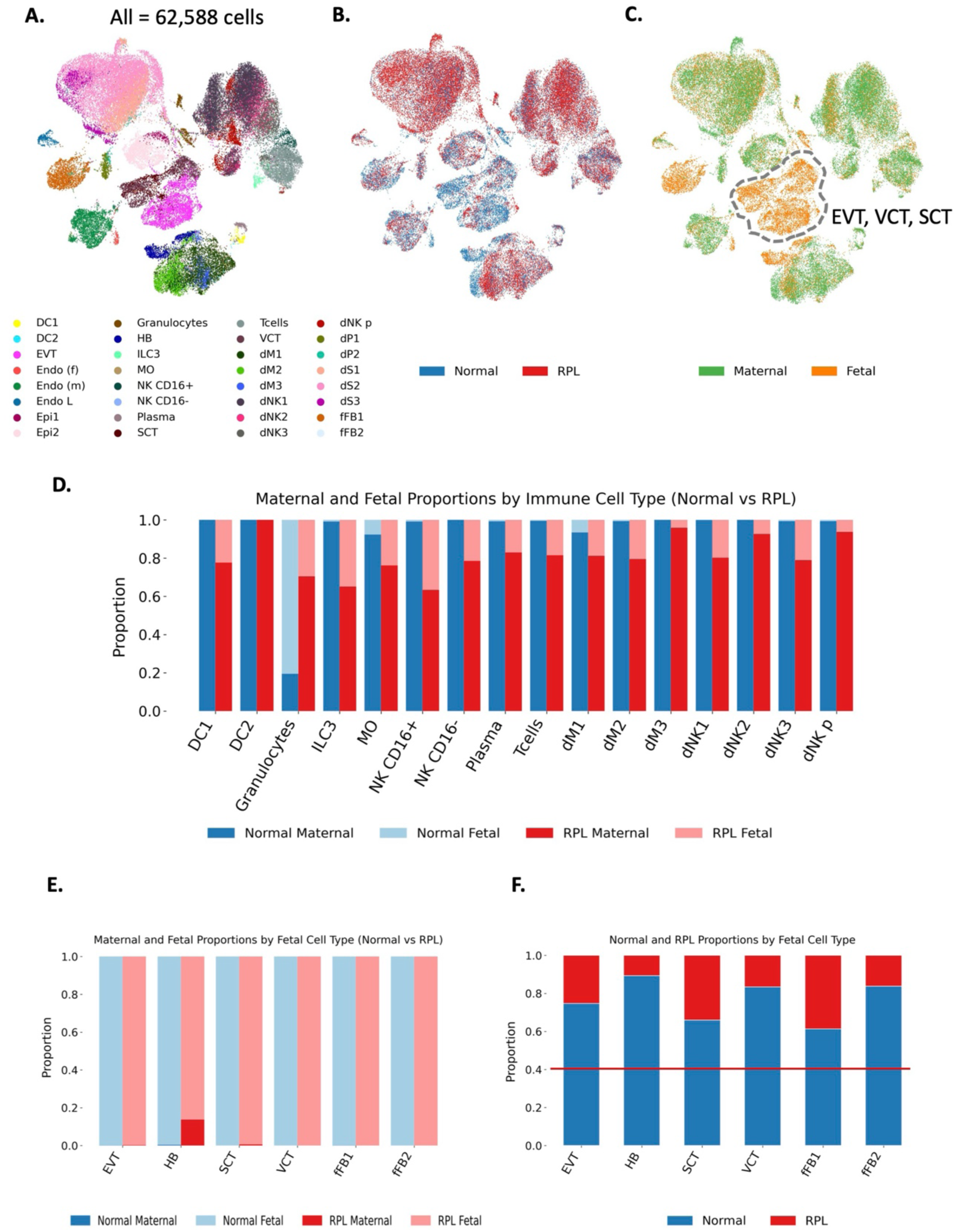
Single-cell landscape of the human decidua from elective terminations of normal pregnancies and RPL. (A) Uniform manifold approximation and projection (UMAP) embedding of cells from our study; each dot represents a single cell and is colored by cell type. Endo, endothelial cells; f, fetal; L, lymphatic; m, maternal; p, proliferative; Epi, epithelial glandular cells; SCT, syncytiotrophoblasts; EVT, extravillous trophoblasts; VCT, villous cytotrophoblasts; DC, dendritic cells; dM, decidual macrophages; dS, decidual stromal cells; HB, Hofbauer cells; ILC, innate lymphoid cells; MO, monocytes; dNK, decidual natural killer cells; dP, decidual progenitor cells. (B) UMAP colored by clinical condition (Norm vs. RPL). (C) UMAP colored by inferred origin (maternal vs. fetal). (D) Stacked bar plots showing the proportions of maternal and fetal cells among immune cell types between Norm and RPL samples. (E) Stacked bar plots showing the proportions of maternal and fetal cells among fetal-derived cell types. (F) Stacked bar plots showing the proportions of cells from Norm and RPL samples among fetal-derived cell types. The red horizontal line indicates the expected baseline proportion of RPL cells across the entire dataset (0.6).

Based on the validated cell origin classification, we next compared the composition of major cell types between the RPL and Norm samples. Immune cell types such as decidual NK cells (dNK), NK CD16+ cells, and decidual macrophages (dM) were primarily maternal in origin (Fig. 1D). Notably, fetal immune cells were nearly absent in Norm samples but were prominently detected in RPL samples. These differences suggest altered immune cell dynamics at the maternal–fetal interface. Conversely, trophoblast lineages and Hofbauer cells—both of fetal origin—exhibited lower proportions of RPL-derived cells than their overall representation in the dataset (0.6), as shown in Fig. 1F. This underrepresentation suggests a selective deficiency of these fetal cells in RPL beyond mere differences in sample size. Considering the critical roles of trophoblasts in placentation and fetal development, this reduction may reflect impaired trophoblast development or differentiation, potentially contributing to insufficient placental formation and compromised fetal support in RPL.

Overall, the single-cell transcriptomic data of our study successfully characterized the cellular composition of the decidua in both Norm and RPL samples. Most immune cell types are predominantly of maternal origin (81% of NK cells across six subtypes and 80% of T cells are maternal), whereas cell types such as trophoblasts are mainly fetal cells (Table S2). In particular, the near absence of fetal immune cells in Norm samples, in contrast to the marked increase in fetal immune cells in RPL samples, indicates a shift in immune cell composition that may reflect altered immune dynamics of fetuses under RPL status. Furthermore, the significantly lower proportion of trophoblast cells in RPL samples suggests that critical trophoblast lineages may fail to develop or differentiate properly, indicating insufficient placental formation. Taken together, these findings suggest that our dataset provides an abundant maternal contribution to understanding RPLs.

### Immune Dysregulation in Euploid RPL Is mutually exclusive to Aneuploid RPL cases

To examine the specificity of the identified immune impairment phenomena in our RPL data, we sought to determine whether disease-associated transcriptional patterns reflect intrinsic features of euploid RPL or overlap with changes observed in RPL accompanied by chromosomal abnormalities (aneuploid RPL). Chromosomal aneuploidy in single cells was inferred using a validated method that integrates the expression of dosage-sensitive genes (via scploid) and allelic imbalance at heterozygous SNP loci. Based on this analysis, fetal-derived decidual cells with chromosomal abnormalities were designated the aneuploid subset and compared against euploid fetal cells from previously karyotyped euploid samples.

We then performed analyses of differentially expressed genes (DEGs) between the RPL and Norm conditions separately within each subset and compared the resulting patterns (Table S3, S4). The analysis first revealed a set of immune-related genes that was consistently upregulated irrespective of ploidy. This group, which includes IL15, IGFBP1, IGFBP2, IGFBP7, and PRDM1, suggests a degree of RPL-intrinsic immune activation.(33–37) In contrast, other gene sets showed starkly opposing regulation depending on ploidy. Several immune-activation genes associated with T/NK cell signaling(38–44) (IL2RB, LCP2, ADGRE5, PTPRJ, PDE7A, RASA2) were upregulated in euploid RPL but downregulated in aneuploid RPL. Conversely, genes related to mitochondrial metabolism and proteostasis(45–50) (MRPL/MRPS subunits, COX5B, NDUFA11, VDAC1, PRDX3, PSMB5, CTSL) exhibited the opposite trend, with strong upregulation in the aneuploid subset. This distinction substantiates our focus on euploid samples and reinforces the biological significance of the transcriptomic signatures identified in later analyses.

### Altered immune response signatures and reduced tolerance in dNK cells of RPL

We hypothesized that RPL arises from a failure of maternal immune tolerance to the fetus followed by rejection. To test this hypothesis, we employed the CRM score, a quantitative measure of the immune response derived from the consistent upregulation of 11 genes during acute rejection in diverse organ transplant settings.^23^ The CRM score was calculated using a gene set scoring approach that compares the expression levels of these CRM genes against a set of reference genes with similar expression distributions (see Methods). A comparison of the CRM scores across all cells revealed that the RPL samples had significantly higher scores compared with the Norm samples. (Mann‒Whitney U test, *p value =* 1.27E-144), suggesting overall activation of immune response pathways in RPL (Fig. 2A).

**Figure 2.**
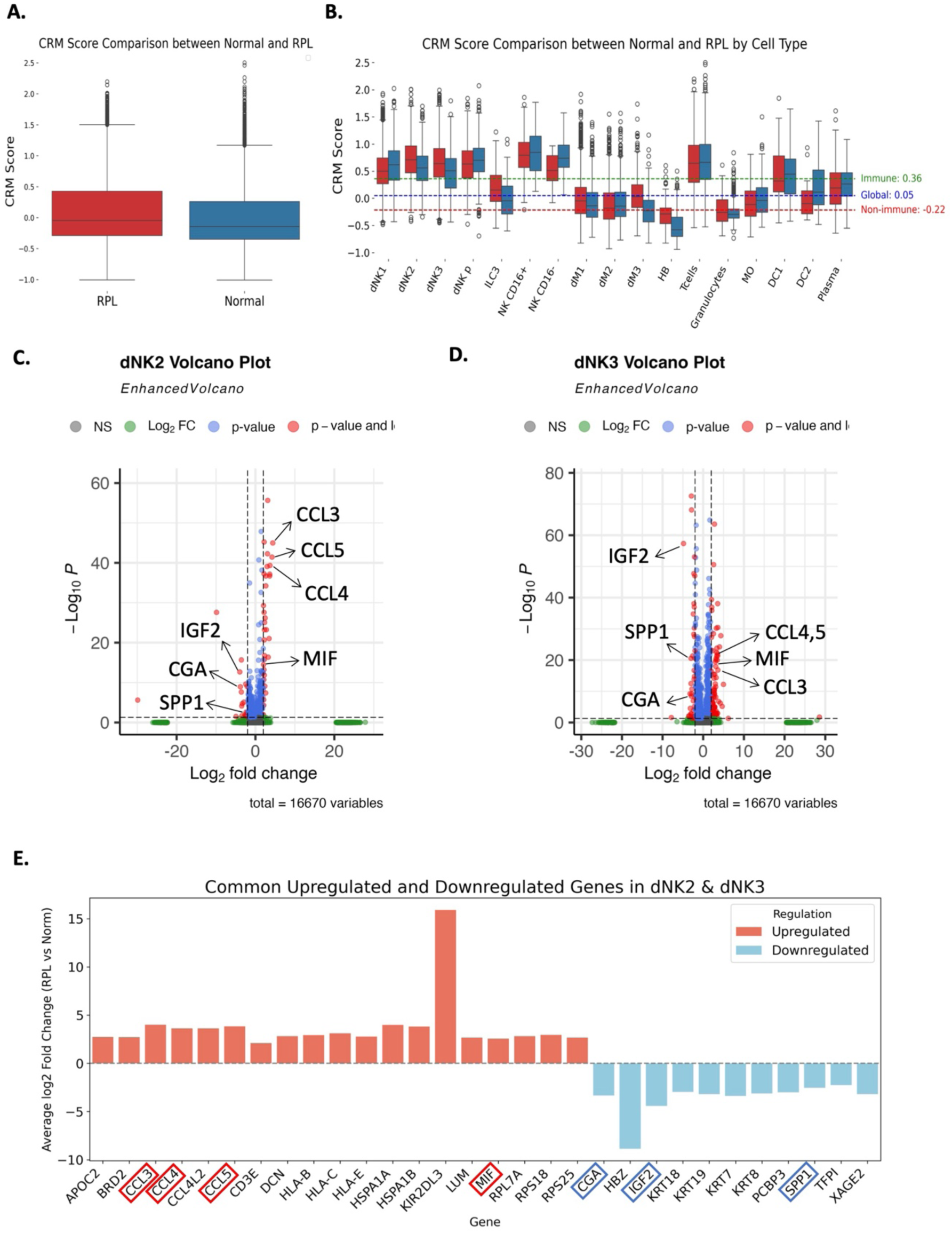
Altered immune response signatures and reduced tolerance in dNK cells from RPL patients. (A) Comparison of common rejection module (CRM) scores between Norm and RPL samples across all cells, showing elevated CRM scores in RPL. (B) Comparison of cell type-specific CRM scores between Norm and RPL samples. Among all immune cell types, dNK2 and dNK3 had the most significantly increased CRM scores in RPL, indicating heightened immune activation. Global, immune, and nonimmune mean CRM score baselines are indicated with dotted lines. (C–D) Volcano plots showing genes that are differentially expressed between RPL and Norm samples in dNK2 (C) and dNK3 (D) cells. DEGs were defined as those whose adjusted p values were < 0.05 and absolute log₂-fold changes were > 2. (E) Bar plot summarizing genes commonly upregulated and downregulated in both the dNK2 and dNK3 subtypes. Proinflammatory chemokines (e.g., CCL3, CCL4, and MIF) were markedly upregulated, whereas immune tolerance–related genes (e.g., IGF2, CGA, and SPP1) were consistently downregulated, indicating a shift toward immune activation and reduced immune tolerance in RPL-associated dNK cells.

To determine which specific immune cell subtypes contributed to the elevated CRM score observed in RPL, we calculated the CRM scores for each cell and compared them across individual cell types. CRM scores were significantly higher in dNK2 (Mann–Whitney U test, *p* = 7.55E-13) and dNK3 (p = 1.23E-17) cells from RPL samples than in those from Norm samples (Fig. 2B). We mark that dNK2 are mainly consist of maternal cells. Under Norm conditions, dNK cell subtypes, particularly dNK2 and dNK3, are critical for supporting pregnancy: dNK2 cells recruit extravillous trophoblasts (EVTs) and dendritic cells to maintain a healthy maternal–fetal interface, and dNK3 cells regulate EVT invasion via CCL5 expression.(51) Moreover, both subtypes actively modulate the local immune microenvironment through the secretion of cytokines and chemokines such as XCL1, CCL3, and CCL4.(52) The observation that these subtypes have significantly higher CRM scores in RPL suggests that their immunomodulatory roles may be altered under pathological conditions. The transcriptional signatures of dNK2 and dNK3 in RPL samples resemble those observed in acute transplant rejection cases, whereas the same cell types in Norm samples do not exhibit this pattern. As shown in Fig. 1D, the majority of these dNKs are maternal cells. Thus, the maternal immune signature contributes primarily to increased CRMs in RPLs.

To investigate the transcriptional alterations underlying the increased immune activation observed in dNK2 and dNK3 cells from RPL samples, we conducted DEG analyses to compare RPL and Norm samples within each subtype (Fig. 2C, 2D; Table S5). In particular, we focused on genes whose expression levels differed by more than fourfold between the two conditions, as such changes are likely to represent biologically meaningful transcriptional shifts. We identified a set of genes that were consistently dysregulated in both dNK2 and dNK3 cells (Fig. 2E). Notably, the expression levels of genes involved in chemokine-mediated immune cell recruitment (CCL3, CCL4, and CCL5) were markedly upregulated in RPL, suggesting enhanced infiltration of T cells, monocytes, and NK cells through CCR5 signaling and a proinflammatory microenvironment at the maternal–fetal interface.(53) The expression of the innate immune regulator MIF was also strongly upregulated, supporting its role in amplifying cytokine production and immune cell activation.(54) Conversely, the expression levels of several key genes related to immune tolerance were downregulated in RPL dNK2 and dNK3 cells (Fig. 2E). The expression of IGF2, a growth factor essential for trophoblast development and immune regulation,(55) was markedly reduced in RPL, which is consistent with its established role in promoting fetal growth and limiting inflammation during pregnancy. CGA, which encodes a subunit of human chorionic gonadotropin (hCG) and contributes to regulatory T-cell expansion while suppressing NK cell cytotoxicity, was also downregulated.(56) Similarly, the expression of SPP1, a gene involved in immune cell adhesion and migration, was reduced.(57) The concurrent upregulation of immune-activating genes and downregulation of tolerance-related genes may collectively contribute to disrupted maternal–fetal immune regulation in RPL.

Overall, our analysis compared the CRM scores across various cell types between the Norm and RPL samples and revealed that dNK2 and dNK3 cells had significantly elevated CRM scores. Differential gene expression analyses of dNK2 and dNK3 cells revealed transcriptional patterns consistent with immune activation and loss of tolerance from maternal cells, supporting our hypothesis that RPL arises from a failure of immune tolerance similar to acute rejection in organ transplantation. However, the imbalance in our dataset poses several limitations for this analysis. For instance, the number of Norm fetal dNK1 cells was extremely low (only 2 out of 8,626 cells), which significantly reduced the statistical power of our comparisons and increased the risk of bias. Such data imbalance may lead to the underestimation or overestimation of true differences between the Norm and RPL groups. These limitations underscore the need for alternative analytical strategies that can effectively overcome data imbalances, thereby enabling the identification of robust transcriptomic signatures and biomarkers related to the failure of maternal–fetal immune tolerance in RPL.

### Machine Learning-based Multilevel Classification Prioritized Transcriptomic Signatures in RPL

Building on the need for an alternative analytical strategy capable of overcoming data imbalance, we adopted a supervised machine learning framework. Instead of testing for mean differences in individual genes, this approach learns the multivariate patterns that best classify cells into RPL or Norm states. By training the model to capture the subtle, combinatorial signatures across the transcriptome, we aimed to identify robust molecular signatures that could reliably distinguish the RPL condition, which might be missed by traditional statistical methods. To effectively disentangle the gene signatures defining cell identity from the more subtle signatures associated with the RPL condition, we employed a stepwise, hierarchical machine learning-based approach. This method was designed to first identify cell type–specific single-cell transcriptomic signatures, and subsequently extract the gene expression patterns specifically associated with RPL within those identified cell types. The architecture of our multilevel classification model, which was implemented using the devCellPy framework and structured into two hierarchical levels, is shown in Fig. 3A. In the first level, the model distinguishes 32 distinct cell types based on trained gene signatures. In the second level, immune cells identified from the first level are further classified into 34 subtypes labeled as either Norm or RPL. For instance, T cells at the first level are reclassified at the second level as T cells from Norm samples or T cells from RPL samples. Although we initially considered implementing a third level of classification to distinguish maternal versus fetal origins among immune cells, this was not feasible because of the extremely small number of fetal-derived immune cells in Norm pregnancy samples. For example, among the 1,404 T cells classified as Norm, only 6 cells (0.04%) were identified as fetal derived, which limited the ability of the model to learn from this class (Table S2).

**Figure 3.**
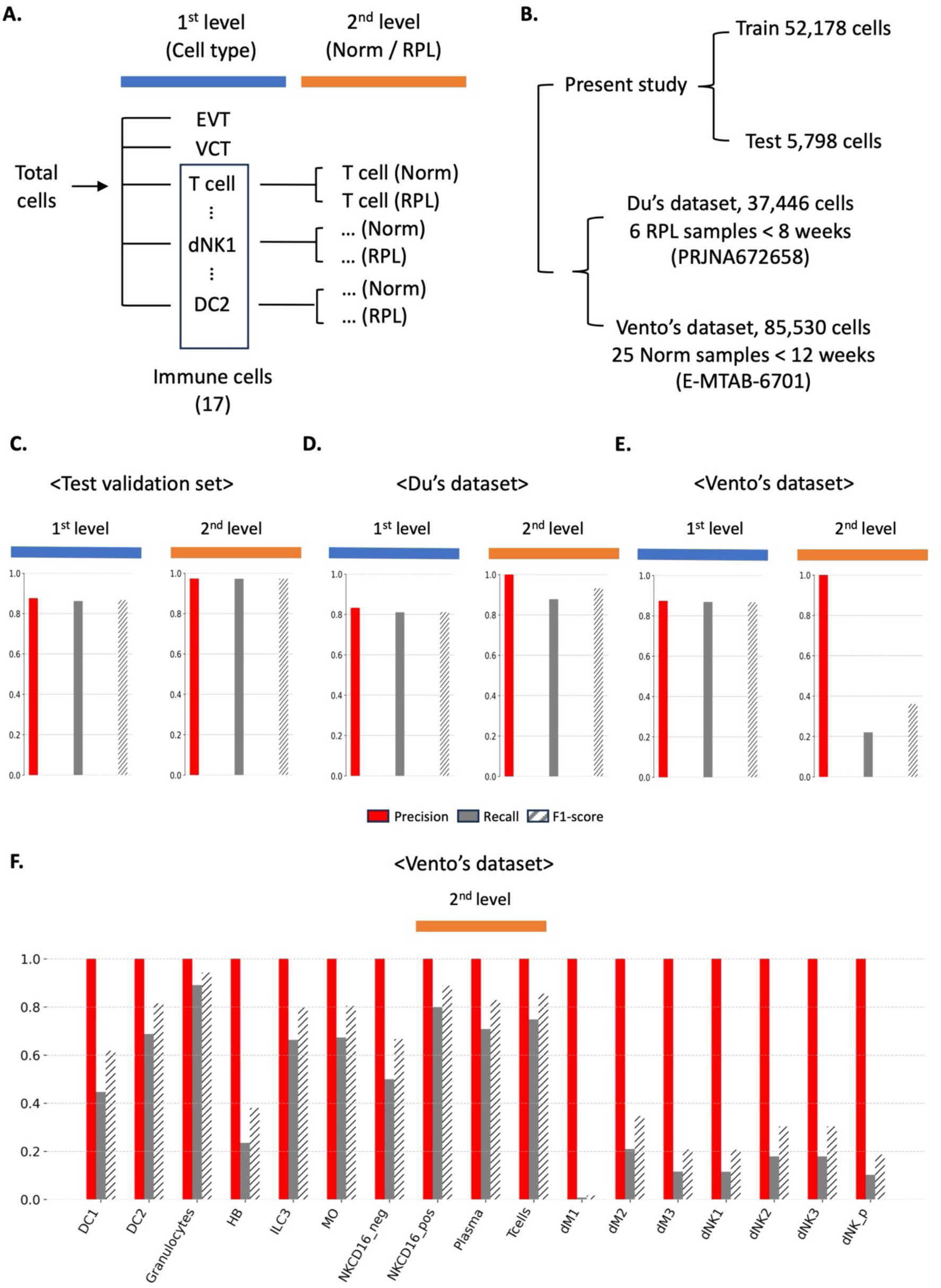
The machine learning approach revealed distinct immune cell signatures linked to RPL. (A) Hierarchy of the classification labels of devCellpy. Using the XGBoost approach, devCellpy allows multilabeled and hierarchically structured identification of cells and assigned subtypes of cells. In the 1st level, the total number of classification labels is 32. Afterward, at the 2nd level, devCellpy is established to identify immune cell types, including Norm (Tcells_Normal) and RPL (Tcells_RPL) T cells, based on pregnancy termination status. (B) Summary of the datasets used in this study. (C) Weighted average precision, recall, and F1 score across all cell labels in the test validation set in the cross-validation settings. The total number of classification labels in each level is also presented. The detailed performance of each cell label at each level is presented in Fig S1. (D-E) Weighted average precision, recall, and F1 score for both 1st- and 2nd-level classifications across all cell labels in the Du and Vento datasets, respectively. (F) Weighted average precision, recall, and F1 score for each cell type at the 2nd level evaluated on Vento’s dataset. The performance at the 1st level is presented in Fig S2.

To assess the robustness of the classification model, we evaluated its performance on a test validation set and two independent validation samples—Du’s dataset and Vento’s dataset (Fig. 3B). Du’s dataset comprised 37,446 cells obtained from decidua samples from six patients (5– 8 weeks gestation; mean 6.83 ± 0.75 weeks), each with ≥ 2 unexplained miscarriages and no fetal chromosomal abnormalities detected by karyotyping or CGH.(58) Vento’s dataset comprised 85,350 cells obtained from decidual tissues from uncomplicated elective terminations between 6–12 weeks of gestation.(7) These datasets were selected to represent distinct biological conditions and sample sources: The test validation set was derived from the same dataset used for model training, whereas the Du and Vento datasets provided independent, external validation from RPL and normal pregnancy contexts, respectively. To ensure this, we tested whether the classifier could maintain predictive accuracy when applied to samples it had never been seen before, including those derived from distinct clinical settings. The trained model was applied to each dataset, and the performance was assessed at both classification levels as described above.

Fig. 3C and 3D depict the classification performance of our multilevel model across each level for the test validation set and Du’s dataset, respectively, while Fig. 3E shows the performance on Vento’s dataset. For the test validation set, the model performed strongly at each level (weighted average F1 score: 0.87 at the first level and 0.97 at the second level). In terms of classification performance by cell type, high performance was observed for the majority of cell types at both levels (Fig S1). In Du’s dataset (Fig. 3D), the classifier achieved a weighted average F1 score of 0.81 at the first level and 0.93 at the second level. As seen in the test validation dataset, classification by cell type also showed high performance for the majority of cell types at both levels (Fig S2). However, in Vento’s dataset, although the model maintained high performance at the first level (weighted average F1 score: 0.87), it underperformed at the second level (weighted average F1 score: 0.32). Nonetheless, the model achieved high classification accuracy in both the training and validation datasets, particularly for immune cell types of interest. Notably, T cells, granulocytes, and NK CD16⁺ cells were consistently classified with high precision in the independent validation set, with weighted average F1 scores of 0.86, 0.94, and 0.89, respectively. These results indicate that these immune cell types possess robust and distinguishable transcriptomic signatures relevant to RPL (Fig. 3F).

In summary, our multilevel machine learning approach combined with SHAP analysis revealed that key immune cell populations, particularly T cells, granulocytes, and NK CD16+ cells, exhibit transcriptomic signatures that distinguish Norm samples from RPL samples. These findings highlight both the common gene contributions across different cell types and unique cell type–specific signals, suggesting that alterations in immune signatures may play a role in the pathogenesis of RPL. Moreover, unlike analyses comparing differences in gene expression between Norm and RPL samples, our approach captures subtle and complex transcriptomic patterns that might otherwise be overlooked, further reinforcing the utility of our method for identifying signature genes associated with RPL.

### Validation of Immune Signatures in RPL using Transformer-based AI

To determine marker genes, which contribute mainly to classification performance at each level, we performed level-specific Shapley additive explanations (SHAP) analysis.(59) We applied SHAP analysis to both the first classification level (cell type distinctions) and the second level (Norm vs. RPL distinctions within immune cells). This approach allowed the quantification of the impact of genes that contributed strongly to the classification at each model layer. For instance, SHAP values were used to rank genes based on their contribution to distinguishing cell types in the first level and T cells from Norm and RPL cells in the second level. In the first-level SHAP analysis, for example, in VCT, a known marker for cytotrophoblasts, such as KRT18, appeared among the top contributors, demonstrating that SHAP reliably identifies label-defining features.(60) This finding demonstrates that SHAP accurately identifies canonical cell-type markers, confirming the validity of our approach and justifying the second-level SHAP analysis to reveal genes that distinguish Norm cells from RPL cells. Table S6 and Table S7 presents the SHAP analysis results for each cell type at each level, including the SHAP values for individual genes.

We independently evaluated the robustness of the SHAP-derived genes and examined whether our devCellPy classifier had overfit by relying on superficial patterns caused by class imbalance. For this, we used scGPT, a transformer-based foundation model trained on large-scale single-cell RNA-seq data.(26) We hypothesized that the top SHAP-ranked genes, identified from the highest-performing immune cell types (T cells, granulocytes, and NK CD16⁺ cells), would contain robust transcriptomic signals that distinguish RPL from Norm. To test this, we fine-tuned scGPT using only these genes and compared its classification performance against a zero-shot baseline, where the model was applied without any fine-tuning. Improved performance in the fine-tuned condition would provide evidence that the selected genes capture biologically meaningful signals that consistently support accurate classification across fundamentally different model architectures and thus serve as distinctive markers of RPL.

The RPL signature was determined by extracting and combining the top 100 SHAP-ranked genes across both classification levels (Table S6, S7) for each of the three immune cell types. We filtered the training dataset to include only the expression values for these genes and modified the model’s input layer accordingly. We then evaluated model performance in two settings: a zero-shot setting using the pretrained scGPT_human model and a fine-tuned setting using our curated training data. In contrast to the two-level devCellPy workflow, the fine-tuned scGPT model assigned six labels in a single step—Tcell_Normal, Tcell_RPL, Granulocyte_Normal, Granulocyte_RPL, NK_CD16⁺_Normal, and NK_CD16⁺_RPL— allowing direct evaluation of the classifier’s ability to simultaneously identify cell identity and RPL status (Fig. 4A).

**Figure 4.**
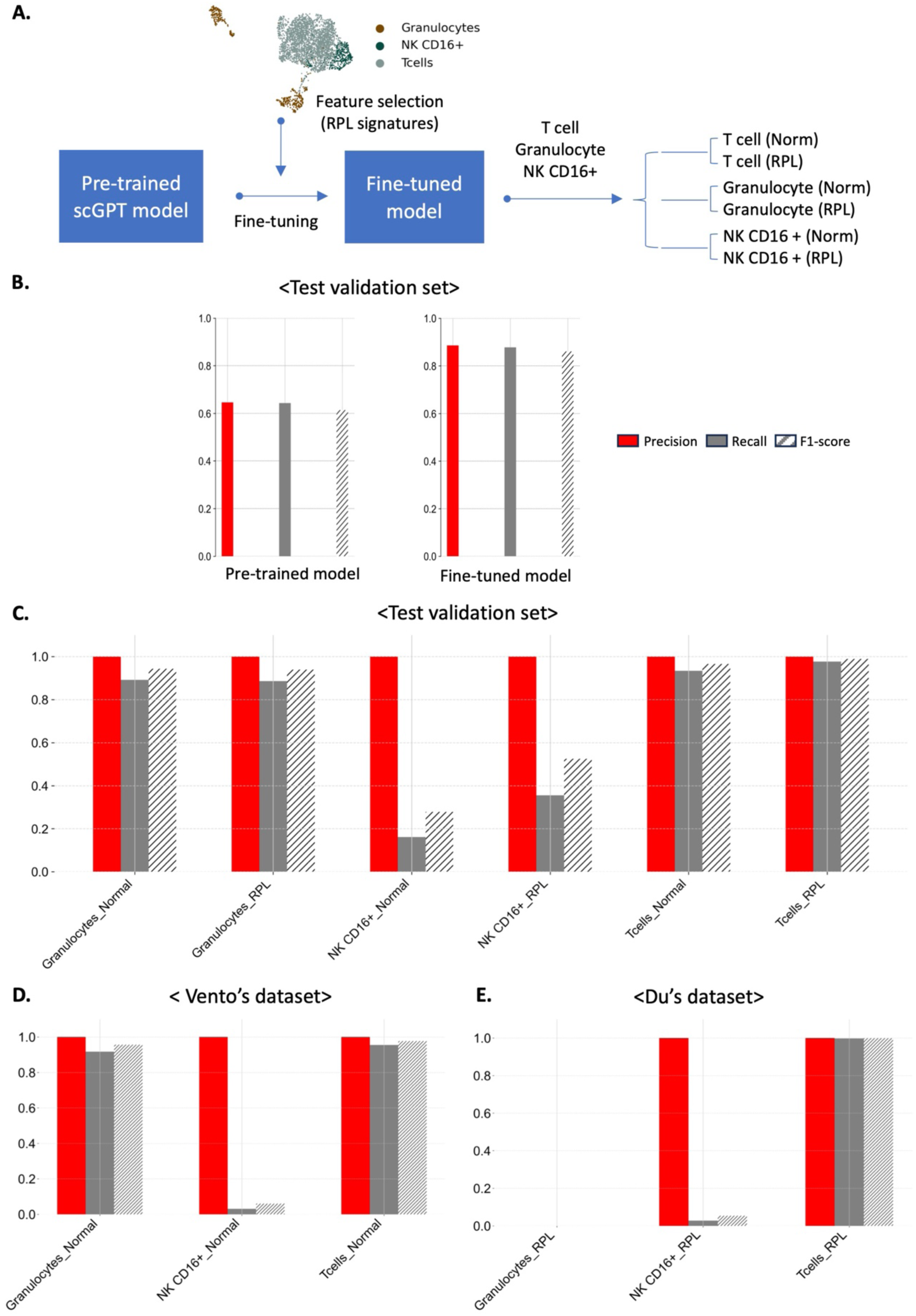
Fine-tuning the scGPT with SHAP-derived gene signatures robustly distinguished the RPL-associated immune cell subtypes. (A) Schematic of the fine-tuning strategy using the transformer-based scGPT_human model. The model was fine-tuned using an RPL gene signature derived from the top SHAP-ranked genes for T cells, granulocytes, and NK CD16⁺ cells. Fine-tuning was performed on expression values restricted to this gene set. (B–E) Model performance was evaluated using the weighted average precision, recall, and F1 score. (B) Comparison of the pretrained and fine-tuned models on the test validation set. (C) Performance of the fine-tuned model on the test validation set for each immune cell subtype. (D) Performance of the fine-tuned model on the independent Norm validation set. (E) Performance of the fine-tuned model on the independent RPL validation set.

As a baseline, before any fine-tuning, the pretrained scGPT_human model achieved a weighted average F1 score of 0.61 on the test validation set. However, our fine-tuned model increased this score to 0.86 (Fig. 4B). This improvement confirms that the selected gene set used for fine-tuning effectively captures the core transcriptomic patterns needed for RPL classification. We then examined the performance by cell type. On the test validation set, the F1 scores of the granulocytes were 0.94 for both Norm and RPL, and the F1 scores of the T cells were 0.97 for Norm and 0.99 for RPL (Fig. 4C). Similarly, the F1 scores of the Vento dataset were 0.96 for granulocytes and 0.98 for T cells (Fig. 4D). We further evaluated the model on the Du dataset. T cells maintained a weighted average F1 score of 0.99, whereas granulocyte performance decreased because of the small number of available cells (n = 9) (Fig. 4E). The observed results demonstrate that the fine-tuned model generalizes well across datasets and cell types, particularly for T cells, which consistently exhibit high predictive accuracy. The consistent performance of T-cell–derived RPL signatures across distinct classification frameworks confirms the predictive power of these signatures and highlights their potential for clinical diagnostic applications in RPL.

### Integrative Pathway and Network Analyses of SHAP-Derived RPL Signatures Reveal Immune-Associated Mechanisms and Therapeutic Targets of RPL

We assessed the potential clinical relevance of SHAP-derived gene signatures associated with RPL. Overrepresentation analysis of these RPL signatures revealed enriched cellular functions, with pathways showing adjusted *P* < 0.05, which were considered significant. The top 20 pathways were ranked by odds ratio, defined as the proportion of query genes mapped to each pathway. This analysis revealed marked enrichment of RPL signatures in five biological categories: translation machinery, mRNA quality control, viral translation, ROBO signaling, and nutrient stress response (Fig. 5A, Table S8).

**Figure 5.**
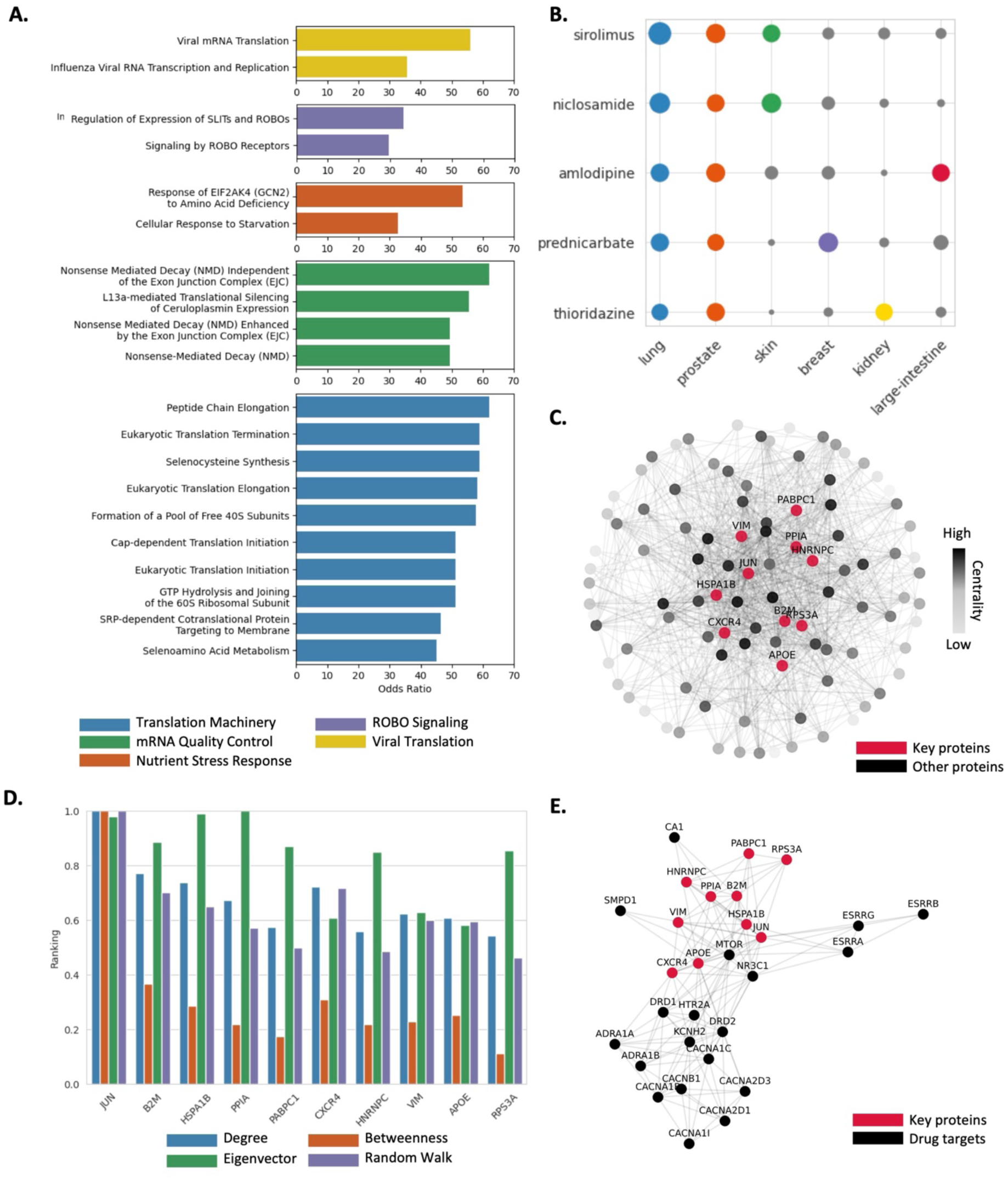
Pathway Analyses and Drug Repurposing Reveal Clinical Implications of T-Cell Signature Genes in RPL. (A) Bar plot showing the results of overrepresentation analysis (ORA) of 100 RPL signatures using the Reactome pathway database. The top 20 pathways with the highest odds ratios, among those with adjusted P values ≤ 0.05, are displayed. Pathways are grouped according to functional categories. (B) Dot plot showing drug repurposing predictions for RPL based on RPL signatures using ASGARD. Each dot represents a predicted drug, with the dot size inversely proportional to the P value. Dots corresponding to drugs with P values ≤ 0.05 are color-coded according to the tissue type in which the prediction was made. (C) Network visualization of interactions between RPL signatures based on STRING protein–protein interaction data. Node color intensity reflects average centrality. The top 10 signature genes with the highest average centrality are labeled and highlighted in red. (D) Bar plot showing the centrality scores of the top 10 signatures in the protein–protein interaction network. Centrality values were min–max normalized to range between 0 and 1. (E) Visualization of the protein–protein interaction network of the top 10 RPL signatures and target proteins of the 5 repurposed drugs.

Among the five functional groups, translation machinery exhibited the greatest number of enriched pathways, which was consistent with the T-cell origin of the RPL signatures. Upon antigen stimulation, T cells exit quiescence and undergo extensive transcriptional and metabolic reprogramming, processes closely linked to translational control during activation and immune regulation.(61–63) The mRNA quality control group was predominantly associated with nonsense-mediated decay (NMD), a post-transcriptional mechanism increasingly recognized for its role in shaping immune responses and emerging as a therapeutic target in autoimmunity.(64,65) ROBO signaling, which is implicated in early trophoblast function, may contribute to RPL through placental dysregulation.(66,67) Although the connections of viral translation and nutrient stress response pathways to immune tolerance are less clearly defined, their relevance is supported by established links between pregnancy loss and immune responses.(68–70) Together, these findings underscore the strong immunological component of the RPL transcriptomic signatures, supporting a mechanistic model of immune-mediated pregnancy loss.

We next asked whether reversing RPL-associated transcriptional changes could mitigate RPL pathology. To address this, we applied ASGARD,(27) a drug-repurposing algorithm that matches disease-associated gene expression profiles with compounds from the L1000 drug-response dataset predicted to counteract those signatures (see Methods). ASGARD identified five candidate compounds predicted to significantly reverse RPL-associated gene expression in at least three distinct tissues (Fig. 5B, Table S9). All five drugs showed predicted efficacy in lung and prostate tissues, with variable activity across other tissues. Notably, three compounds (prednicarbate, sirolimus, and niclosamide) have previously been implicated in contexts relevant to RPL, providing orthogonal support for their therapeutic potential.

Corticosteroids such as prednicarbate are typically used to treat autoimmune conditions and are administered to women with repeated early pregnancy loss.(71,72) Recent meta-analyses report a significant increase in ongoing pregnancy rates following corticosteroid treatment.(73) Sirolimus (rapamycin), which inhibits T- and B-cell activation and proliferation, is approved for use in transplantation and oncology.(74,75) A recent phase II randomized trial revealed that sirolimus significantly improved both pregnancy and live birth rates in patients with repeated implantation failure.(76) Niclosamide has been proposed as a therapy for immune-mediated disorders, including autoimmune diseases and infections, through the modulation of immune pathways.(77) Notably, its inhibition of follicular helper T cells has been shown to be effective in preclinical autoimmune models.(78) Although neither thioridazine nor amlodipine has been directly associated with RPL or immune tolerance, calcium channel blockers such as amlodipine exhibit known immunosuppressive effects that may warrant further investigation.(79,80) These findings support the validity of our drug-repurposing predictions and further implicate RPL gene signatures in the pathophysiology of RPL.

To identify candidate therapeutic targets from the RPL gene signatures, we constructed a protein–protein interaction (PPI) network by mapping RPL-associated genes to their encoded proteins and assembling them using STRING (Fig. 5C). In such networks, hub proteins with high connectivity are more likely to modulate overall network behavior.(81,82) To prioritize central nodes, we computed four network measures (degree centrality, betweenness centrality, eigenvector centrality, and random walk) and ranked the proteins based on their average centrality scores. The top ten hub proteins were selected from the 100 RPL-associated proteins (Fig. 5D, Table S10). These hub proteins exhibited dense connectivity with targets of repurposed drugs identified earlier. In particular, the strong connectivity with mTOR is noteworthy, given its established role in T-cell–mediated immunity and its therapeutic relevance in autoimmune disease contexts (Fig. 5E).

A focused literature review revealed that seven of the ten top-ranked hub proteins are associated with RPL or immune regulation, further supporting their therapeutic potential. CXCR4, for instance, plays a critical role in coordinating both innate and adaptive immune responses.(83) Additionally, a recent systematic review and meta-analysis revealed a significant association between maternal APOE genotype and RPL risk in Asian populations.(84) Single-nucleotide polymorphism analysis involving more than 200 patients further suggested that APOE variants may predispose individuals to RPL.(85) Although JUN, HSPA1B, PPIA, and VIM have not been directly linked to RPL, these genes are functionally implicated in T-cell activity and activation and are known to contribute to immune modulation.(86–92) Together, these findings highlight the functional relevance of the identified RPL signature genes in the pathophysiology of RPL and support their potential as biomarkers and therapeutic targets for future translational studies.

### Confounding-Controlled Expression Filtering Identifies Robust T-cell–associated Transcriptomic Signatures in RPL

Building upon the insights from the functional and network-based characterization of SHAP-derived T-cell signatures, we next sought to further refine the top-ranked genes from the T-cell classifier by evaluating whether their expression patterns remained consistent after accounting for potential confounding effects, such as differences in cell origin or cellular composition. To this end, we performed an origin-controlled expression filtering analysis aimed at identifying genes that exhibit disease-specific transcriptional changes independent of sample-level heterogeneity.

In particular, nearly all the Norm T cells were maternal in origin, whereas a substantial proportion of the RPL T cells were fetal in origin (Table S2). This imbalance raised the possibility that some SHAP-ranked genes might reflect origin-related differences rather than RPL-associated changes. To eliminate this confounding effect, we implemented a two-step validation approach (Fig. 6A). First, we excluded genes whose expression levels significantly differed between maternal and fetal T cells. Next, we retained only those genes that remained significantly differentially expressed between RPL and normal T cells within the maternal subset.

**Figure 6.**
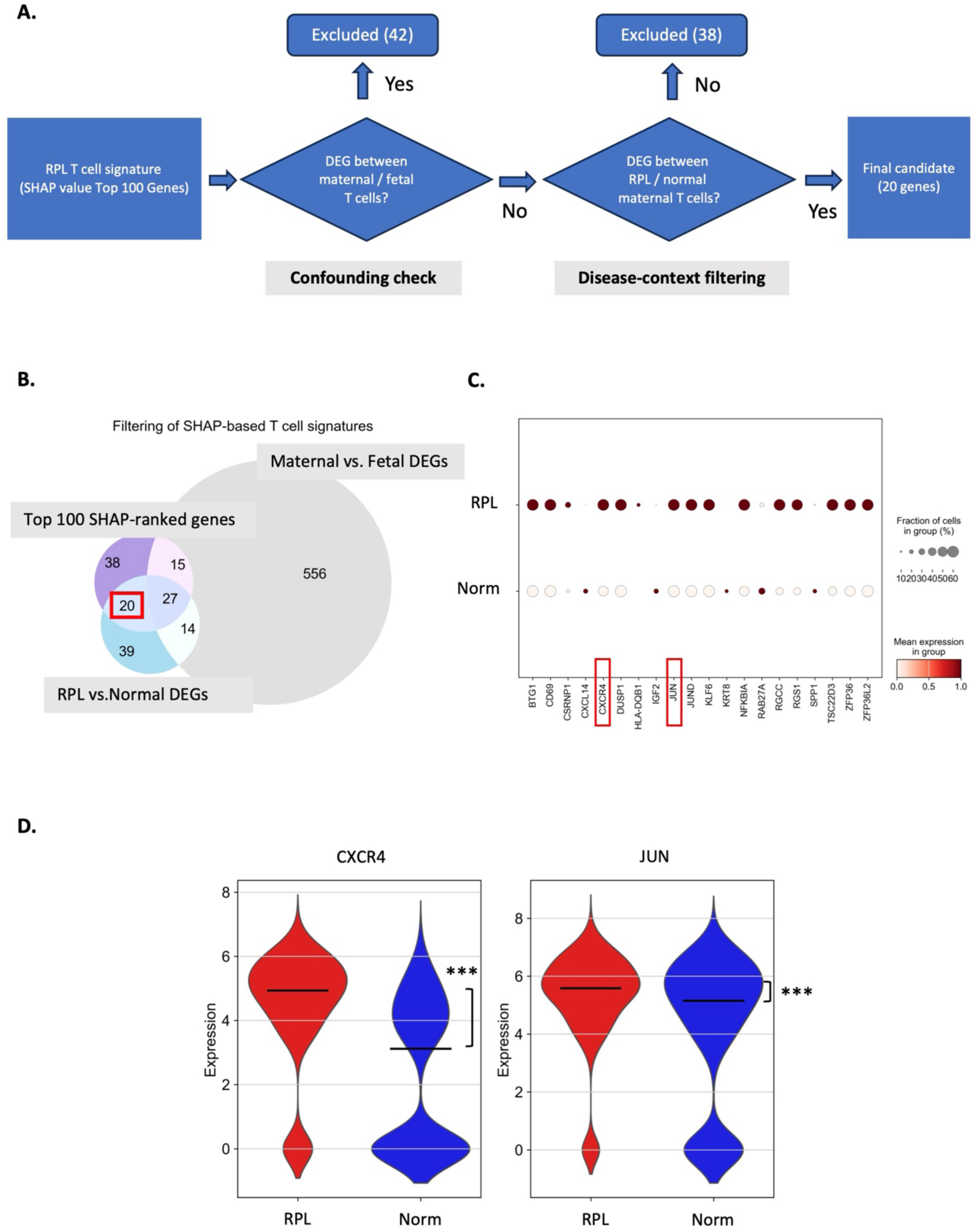
Confounding-controlled filtering identifies robust T-cell signatures in RPL. (A) Schematic of the two-step filtering strategy used to refine SHAP-ranked T-cell genes by removing potential confounding effects. Genes differentially expressed between maternal and fetal T cells (n = 42) were excluded to control for cell origin-associated bias (confounding check). Among the remaining genes, those that were not significantly differentially expressed between RPL and Norm maternal T cells (n = 38) were further excluded. This yielded a final panel of 20 genes showing RPL-specific expression patterns. (B) Venn diagram illustrating the overlap among the top 100 SHAP-ranked genes, maternal vs. fetal T-cell DEGs, and RPL vs. Norm maternal T-cell DEGs. The red-boxed sector (n = 20) represents genes included in the SHAP top set and RPL-related DEGs but not in maternal–fetal DEGs—comprising the final RPL-specific candidates. (C) Dot plot showing the expression and cellular prevalence of the 20 final candidate genes in RPL (RPL_chrNorm) and Norm (normTerm) T cells. The dot size indicates the fraction of cells expressing the gene; the color indicates the average expression. (D) Violin plots of CXCR4 and JUN expression in RPL vs. Norm maternal T cells. Both genes were retained after filtering and showed elevated expression in RPL-T cells (*** indicates p < 0.001 according to the Wilcoxon rank-sum test).

This filtering process resulted in a panel of 20 genes (Fig. 6B), with 13 showing elevated expression in RPL and widespread cellular distribution (Fig. 6C). These genes represent strong candidates for RPL-specific immune markers, supported by both their machine learning–based prioritization and disease-contextual expression validation.

Notably, after origin-controlled filtering, CXCR4 and JUN were the two genes among the 20 genes retained that overlapped with the top hub genes identified in the previous network-based analysis. This convergence between independently derived gene sets further supports their biological relevance and highlights them as particularly robust candidates implicated in RPL. The expression levels of both genes were significantly greater in RPL T cells than in Norm cells (Fig. 6D).

CXCR4 encodes a chemokine receptor that regulates leukocyte trafficking and inflammatory responses, particularly through the CXCL12–CXCR4 axis.(93) In the context of normal pregnancy, CXCR4 has been implicated in promoting immune tolerance and tissue remodeling, including decidual NK cell migration and epithelial repair.(94,95) However, dysregulated CXCR4 expression has been linked to excessive T-cell infiltration and sustained inflammation in various disease settings, such as autoimmunity and cancer.(96,97)

Another gene of interest was JUN, a key component of the AP-1 transcription factor complex that integrates TCR and costimulatory signals to activate T-cell gene programs.(98,99) Upon stimulation, JUN translocates to the nucleus and initiates chromatin remodeling and cytokine transcription.(100,101) Although the role of JUN in normal pregnancy is less well characterized, it has been implicated in cytokine regulation and trophoblast invasion—critical processes in early gestation.(102) In pathological contexts, JUN overexpression in T cells can sustain inflammatory responses and prevent exhaustion, thereby promoting persistent immune activation.(87,103)

Together, these findings suggest that CXCR4 and JUN may contribute to pathological immune activation at the maternal–fetal interface in RPL. To our knowledge, neither gene has been previously highlighted as a distinguishing feature of decidual T cells in the context of normal pregnancy or RPL, underscoring their potential novelty and relevance as immune signatures in pregnancy loss.

## Discussion

In this study, we employed single-cell RNA sequencing and advanced computational modeling to elucidate how immune tolerance fails in RPL. To ensure accurate cellular classification, we incorporated genotype validation techniques, including Souporcell, which allowed us to distinguish maternal cells from fetal cells reliably. This distinction enabled a detailed understanding of the respective roles these cell populations play in the pathophysiology of RPL. By comparing decidual tissues from Norm and RPL cases, our cell origin analysis revealed that most immune cells were predominantly maternal, whereas trophoblast lineages such as EVT and SCT were overwhelmingly fetal in origin. In the RPL samples, the number of fetal-derived trophoblast cells were markedly decreased, whereas the number of fetal immune cells, which are rarely detected in normal samples, were substantially increased. These results highlight the unusual expansion of fetal immune cells and impaired placental development associated with RPLs. Interestingly, the CRM score, a metric developed to quantify rejection responses in organ transplants, of RPL cells resulted in elevated CRM scores compared with those of Norm cells, which were primarily driven by dNK2 and dNK3 maternal cells, supporting our hypothesis; failure of maternal immune harmonization would prompt a proinflammatory state, as in organ rejection. Thus, in RPL without aneuploidy, we pinned decidual NK cells as contributors to pathological conditions. Using the machine learning framework devCellPy, we identified T-cell gene signatures that differentiate RPL from Norm conditions and validated them using the transformer-based model scGPT, confirming their robustness across distinct modeling approaches. Pathway, network, and in silico drug repurposing analyses prioritized immune-associated hub genes and three clinically relevant drug candidates with established safety profiles. Finally, origin-controlled filtering removed cell origin–related confounding, with CXCR4 and JUN emerging as robust candidates shared between filtered RPL-specific T-cell signatures and network-based hub genes. Taken together, our integration of dual confirmation– based origin assignment with complementary AI models represents a methodological advance broadly applicable for uncovering cell type–specific signatures in diverse biological contexts.

Despite the insights gained, several limitations should be noted. First, our study relied on a relatively small and clinically heterogeneous cohort (Norm n = 2, RPL n = 3), which may limit the statistical power to generalize the findings, particularly for rare cell populations and individual-specific transcriptional patterns. However, our analysis focused mainly on transcriptional differences for each cell rather than germline variants; thus, our biological replicates were counted according to the number of cells (62,588 cells). Moreover, we attempted to mitigate this limitation through validation across multiple independent datasets encompassing a broader range of biological contexts. Second, because the CRM score was originally developed to quantify immune responses in transplant settings, its application to maternal–fetal immune harmonizations may require cautious interpretation. In addition, we acknowledge that the small fraction of fetal T cells in the RPL samples prevented the application of our systemic approach. For the same reason, direct identification of the interactions between fetal and maternal immune cells is constrained by the proportional differences in cell origins. Although CXCR4 and JUN have emerged as robust RPL-associated immune signatures, our findings do not establish a causal role for these genes in the pathogenesis of RPL. Given their broader involvement in immune regulation, it remains unclear whether their elevated expression reflects a driving mechanism of pregnancy loss or a secondary response to pathological immune activation. Future studies incorporating functional perturbation and in vivo modeling will be essential to clarify the mechanistic roles of key immune signatures in RPL.

## Conclusion

In conclusion, we investigated the immunological mechanisms underlying RPL at the single-cell level. By integrating single-cell transcriptomics with machine learning and transformer-based deep learning models, we identified robust immune-related gene signatures— particularly in maternal T cells—that distinguish RPL pregnancies from normal pregnancies. Furthermore, network-based prioritization and in silico drug repurposing highlighted clinically relevant pathways and therapeutic candidates, reinforcing the translational potential of our findings. These results lay the groundwork for the future development of noninvasive biomarkers and targeted interventions aimed at improving the diagnosis and stratification of RPL.

## Data availability

The scRNA-seq data generated in this study is available in GEO under the accession number GSE 306259 (https://www.ncbi.nlm.nih.gov/geo/query/acc.cgi?acc=GSE306259). This paper does not report original code for the algorithm. Other source codes for figure vignette and genotype analysis codes from the results of Souporcell are presented (https://github.com/hypaik/PePT_vignette). Any additional information required to reanalyze the data reported in this paper is available from the lead contact upon request.

## Acknowledgments

Not applicable.

## Author information

Tae Lyun Ko and Jaesub Park contributed equally to this work.

## Authors and Affiliations

Center for Biomedical Computing, Korea Institute of Science and Technology Information (KISTI), Daejeon, 34141, Republic of Korea

Tae Lyun Ko & Hyojung Paik

Department of Data and HPC Science, Korea National University of Science and Technology (UST), Daejeon, 34141, Republic of Korea

Tae Lyun Ko & Hyojung Paik

Department of Bio and Brain Engineering, Korea Advanced Institute of Science and Technology, Daejeon 34141, Republic of Korea

Jaesub Park

Cambridge Stem Cell Institute, University of Cambridge, Cambridge, UK

Jaesub Park

Department of Obstetrics and Gynecology, College of Medicine, Myunggok Medical Research Institute, Konyang University, Daejeon 158, Republic of Korea

Jae won Han, Jin Sol Park & Sung Ki Lee

Department of Biological Sciences, Sungkyunkwan University, Suwon, 16419, Republic of Korea

Dongju Leem & Junho Kim

## Contributions

T.L.K. conceived the main idea, acquired data, performed the bioinformatics analysis, prepared all figures and supporting figures, and wrote the draft. J.P. supported the bioinformatics and drug repositioning analyses, prepared the main figures, and wrote the manuscript. D.L. and J.K. supported the bioinformatics analysis. J.W.H. and J.S.P. supported the clinical background section and participated in the revision of the manuscript. S.K.L. participated in the conceptual development of the research, provided the tissue samples used in the study, and contributed to the revision of the manuscript and funding acquisition. H.P. conceived the main idea, designed the bioinformatics analysis, supervised the overall project, and acquired funding. All authors participated in discussion of the results, read and approved the final manuscript.

## Corresponding authors

Correspondence to Hyojung Paik or Sung Ki Lee

## Ethics approval and consent to participate

All participants provided written informed consent. The study was approved by the Institutional Review Board (IRB) of Konyang University Hospital (IRB approval number: KYUH 2022-08-009-005).

## Consent for publication

Not applicable.

