## Supplementary Figures S1-S2 for "Machine learning–driven decoding of maternal immune signatures in repeated pregnancy loss"

Fig. S1

<Test validation set>

**A.**

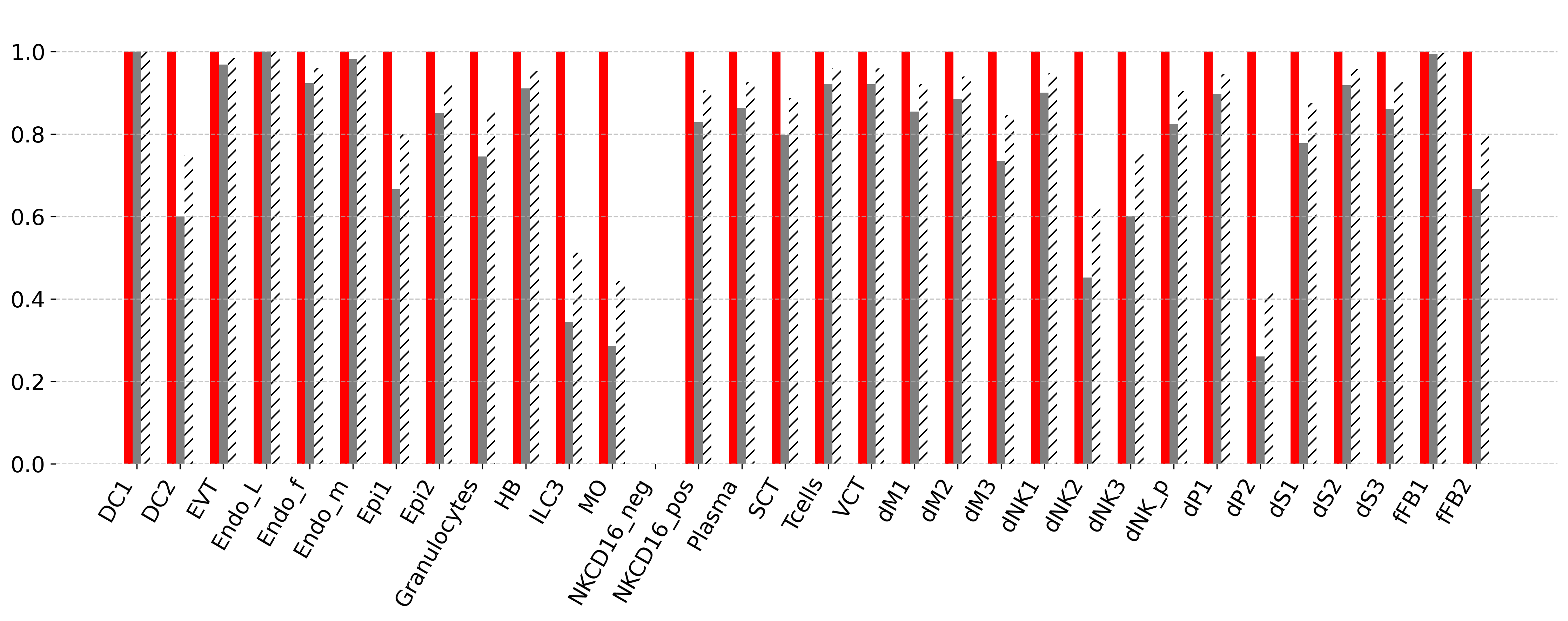

1^st^ level (32 labels)

<Test validation set>

**B.**

2^nd^ level (34 labels)

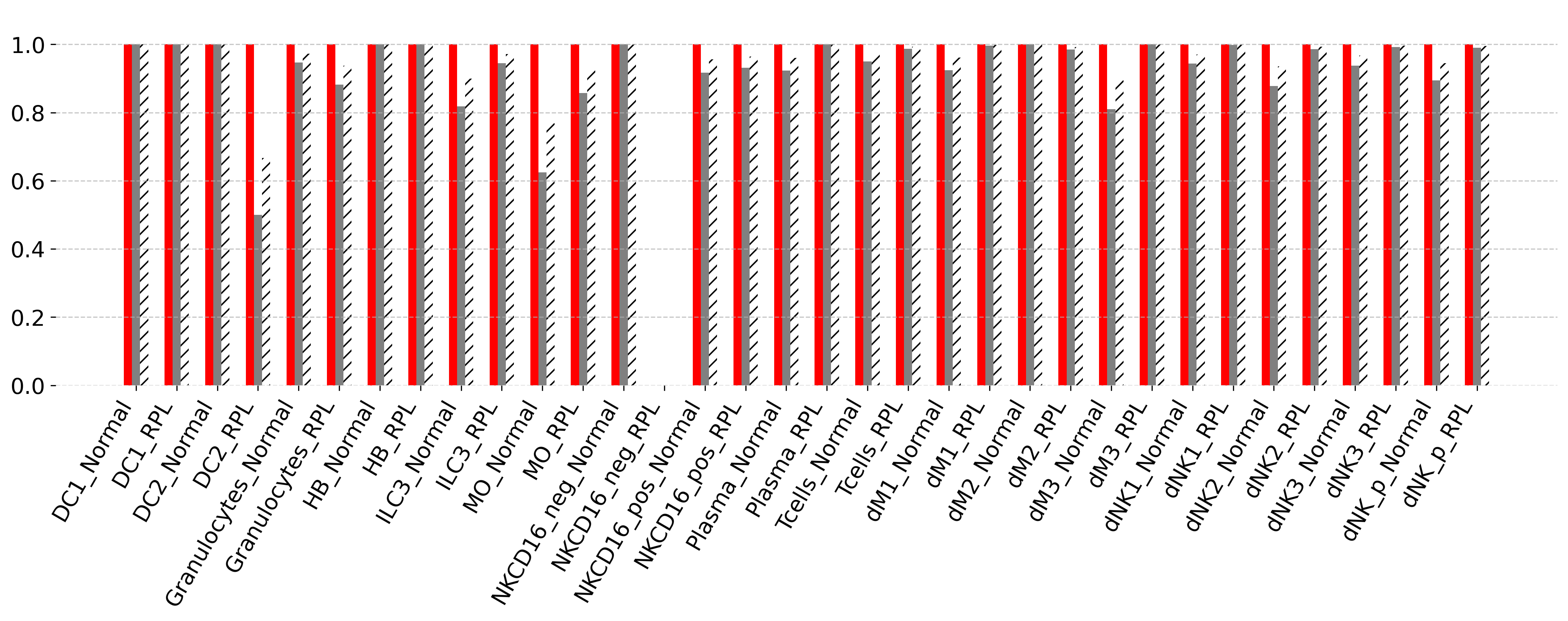

Fig. S1. Detailed performance of the classification model in the test sets

(A–B) The performance of the classification model was evaluated for each cell type in the test validation set, with weighted precision, recall, and F1 score shown for the 1st (A) and the 2nd (B) levels.

Fig. S2

<Du’s dataset>

**A.**

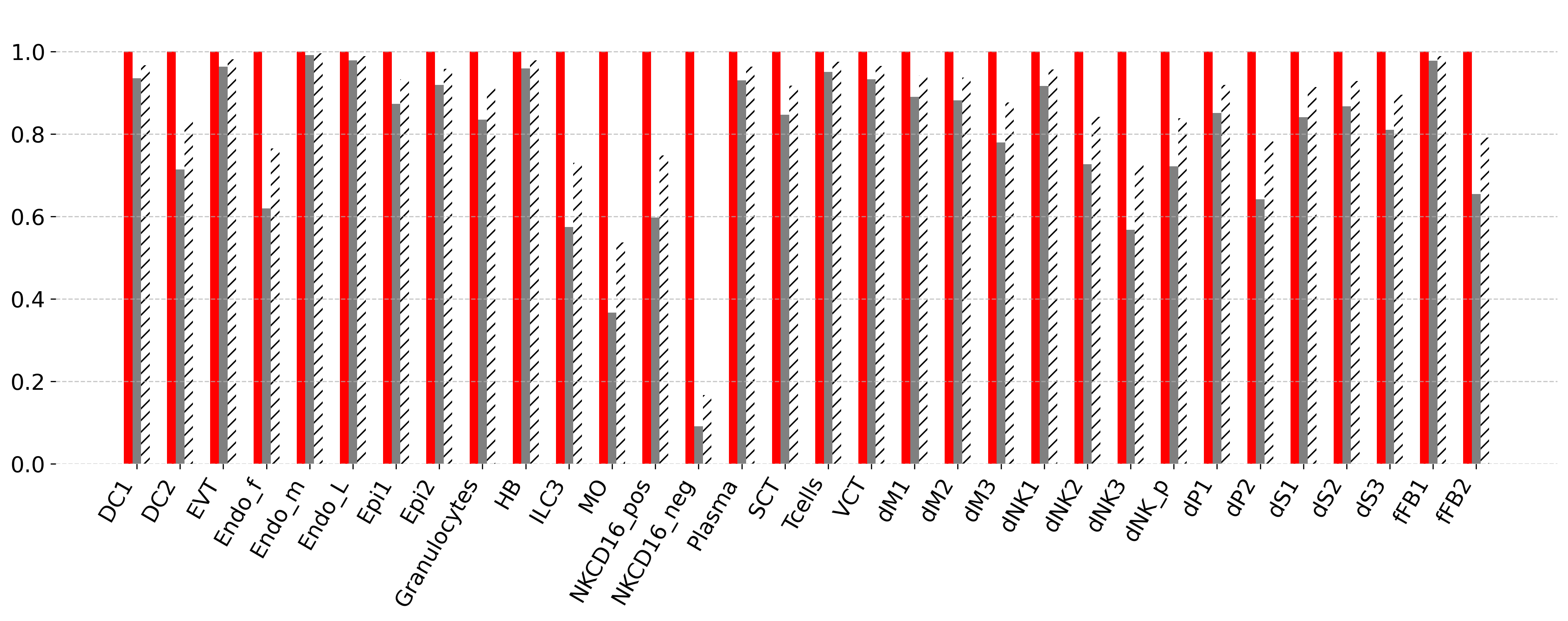

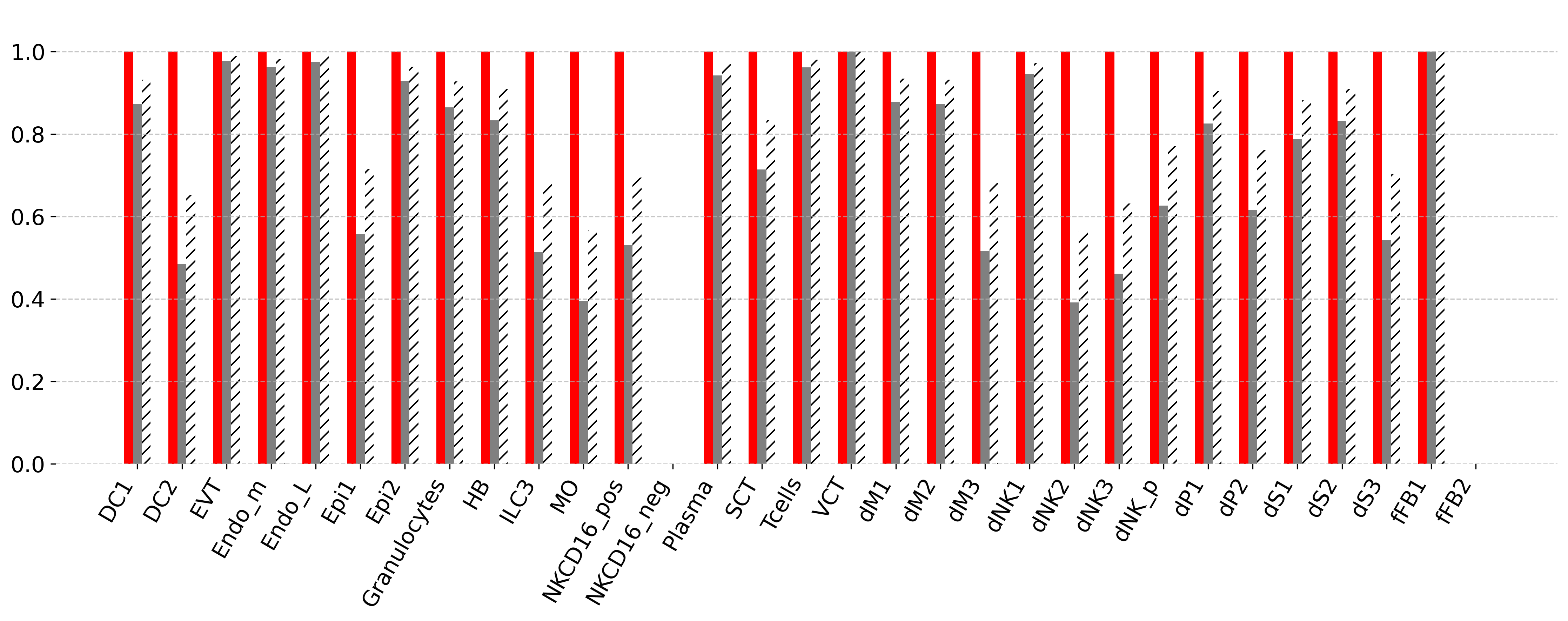

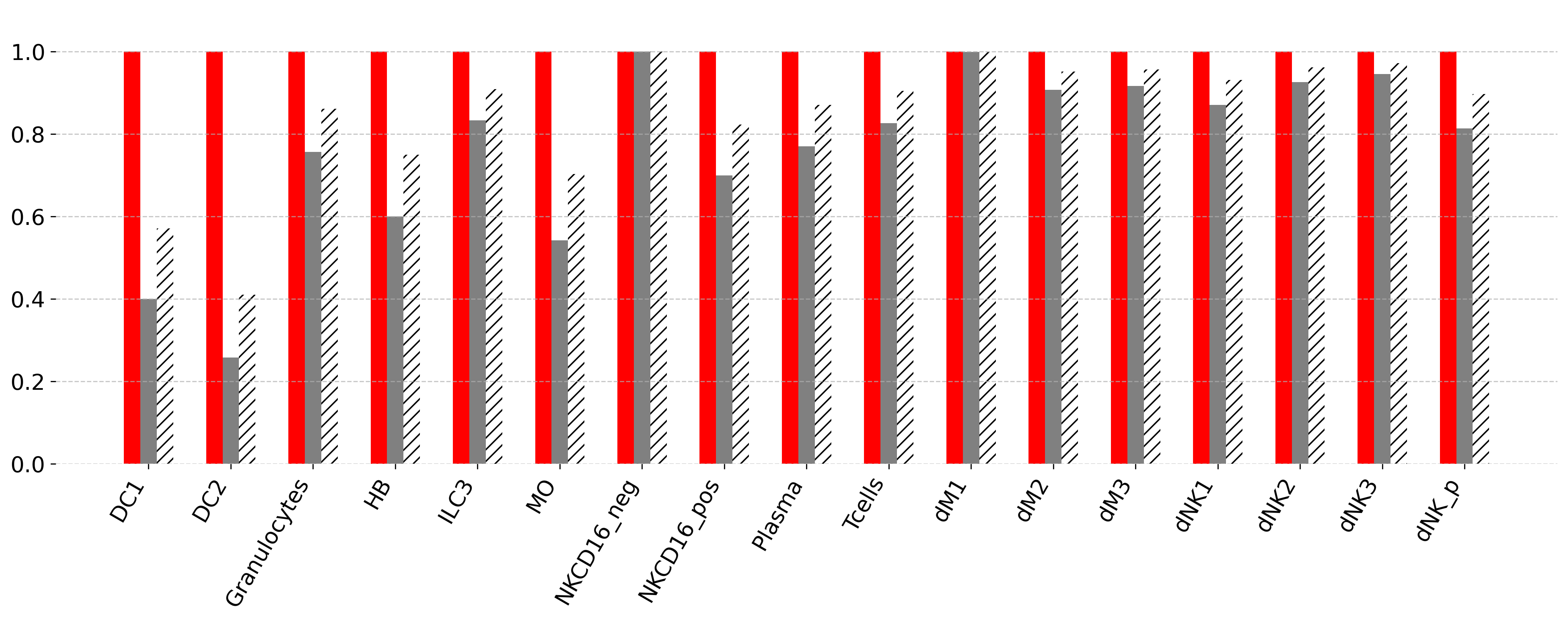

**C.**

1^st^ level

2^nd^ level

<Du’s dataset>

1^st^ level

**B.**

<Vento’s dataset>

Fig. S2. Detailed performance of our established classification model in the validation sets

(A–B) The performance of the classification model was evaluated for each cell type in Du’s dataset, with weighted precision, recall, and F1 score shown for the 1st (A) and 2nd (B) levels. (C) The weighted average precision, recall, and F1 score for each cell type at the 1st level evaluated on Vento’s dataset.
